# Early Pleistocene enamel proteome sequences from Dmanisi resolve *Stephanorhinus* phylogeny

**DOI:** 10.1101/407692

**Authors:** Enrico Cappellini, Frido Welker, Luca Pandolfi, Jazmin Ramos Madrigal, Anna K. Fotakis, David Lyon, Victor J. Moreno Mayar, Maia Bukhsianidze, Rosa Rakownikow Jersie-Christensen, Meaghan Mackie, Aurélien Ginolhac, Reid Ferring, Martha Tappen, Eleftheria Palkopoulou, Diana Samodova, Patrick L. Rüther, Marc R. Dickinson, Tom Stafford, Yvonne L. Chan, Anders Götherström, Senthilvel KSS Nathan, Peter D. Heintzman, Joshua D. Kapp, Irina Kirillova, Yoshan Moodley, Jordi Agusti, Ralf-Dietrich Kahlke, Gocha Kiladze, Bienvenido Martínez-Navarro, Shanlin Liu, Marcela Sandoval Velasco, Mikkel-Holger S. Sinding, Christian D. Kelstrup, Morten E. Allentoft, Anders Krogh, Ludovic Orlando, Kirsty Penkman, Beth Shapiro, Lorenzo Rook, Love Dalén, M. Thomas P. Gilbert, Jesper V. Olsen, David Lordkipanidze, Eske Willerslev

## Abstract

Ancient DNA (aDNA) sequencing has enabled unprecedented reconstruction of speciation, migration, and admixture events for extinct taxa^1^. Outside the permafrost, however, irreversible aDNA post-mortem degradation^2^ has so far limited aDNA recovery within the ˜0.5 million years (Ma) time range^3^. Tandem mass spectrometry (MS)-based collagen type I (COL1) sequencing provides direct access to older biomolecular information^4^, though with limited phylogenetic use. In the absence of molecular evidence, the speciation of several Early and Middle Pleistocene extinct species remain contentious. In this study, we address the phylogenetic relationships of the Eurasian Pleistocene Rhinocerotidae^5-7^ using ˜1.77 million years (Ma) old dental enamel proteome sequences of a *Stephanorhinus* specimen from the Dmanisi archaeological site in Georgia (South Caucasus)^8^. Molecular phylogenetic analyses place the Dmanisi *Stephanorhinus* as a sister group to the woolly (*Coelodonta antiquitatis*) and Merck’s rhinoceros (*S. kirchbergensis*) clade. We show that *Coelodonta* evolved from an early *Stephanorhinus* lineage and that this genus includes at least two distinct evolutionary lines. As such, the genus *Stephanorhinus* is currently paraphyletic and its systematic revision is therefore needed. We demonstrate that Early Pleistocene dental enamel proteome sequencing overcomes the limits of ancient collagen- and aDNA-based phylogenetic inference, and also provides additional information about the sex and taxonomic assignment of the specimens analysed. Dental enamel, the hardest tissue in vertebrates, is highly abundant in the fossil record. Our findings reveal that palaeoproteomic investigation of this material can push biomolecular investigation further back into the Early Pleistocene.

## MAIN TEXT

Phylogenetic placement of extinct species increasingly relies on aDNA sequencing. Relentless efforts to improve the molecular tools underlying aDNA recovery have enabled the reconstruction of ˜0.4 Ma and ˜0.7 Ma old DNA sequences from temperate deposits^9^ and subpolar regions^10^ respectively. However, no aDNA data have so far been generated from species that became extinct beyond this time range. In contrast, ancient proteins represent a more durable source of genetic information, reported to survive, in eggshell, up to 3.8 Ma^11^. Ancient protein sequences can carry taxonomic and phylogenetic information useful to trace the evolutionary relationships between extant and extinct species^12,13^. However, so far, the recovery of ancient mammal proteins from sites too old or too warm to be compatible with aDNA preservation is mostly limited to collagen type I (COL1). Being highly conserved^14^, this protein is not an ideal marker. For example, regardless of endogeneity^15^, collagen-based phylogenetic placement of Dinosauria in relation to extant Aves appears to be unstable^16^. This suggests the exclusive use of COL1 in deep-time phylogenetics is constraining. Here, we aimed at overcoming these limitations by testing whether dental enamel, the hardest tissue in vertebrates^17^, can better preserve a richer set of ancient protein residues. This material, very abundant in the fossil record, would provide unprecedented access to biomolecular and phylogenetic data from Early Pleistocene animal remains.

Dated to ˜1.77 Ma by a combination of Ar/Ar dating, paleomagnetism and biozonation^18,19^, the archaeological site of Dmanisi (Georgia, South Caucasus; Fig 1a) represents a context currently considered outside the scope of aDNA recovery. This site has been excavated since 1983, resulting in the discovery, along with stone tools and contemporaneous fauna, of almost one hundred hominin fossils, including five skulls representing the *georgicus* paleodeme within *Homo erectus*^8^. These are the earliest fossils of the first *Homo* species leaving Africa.

**Figure 1.**
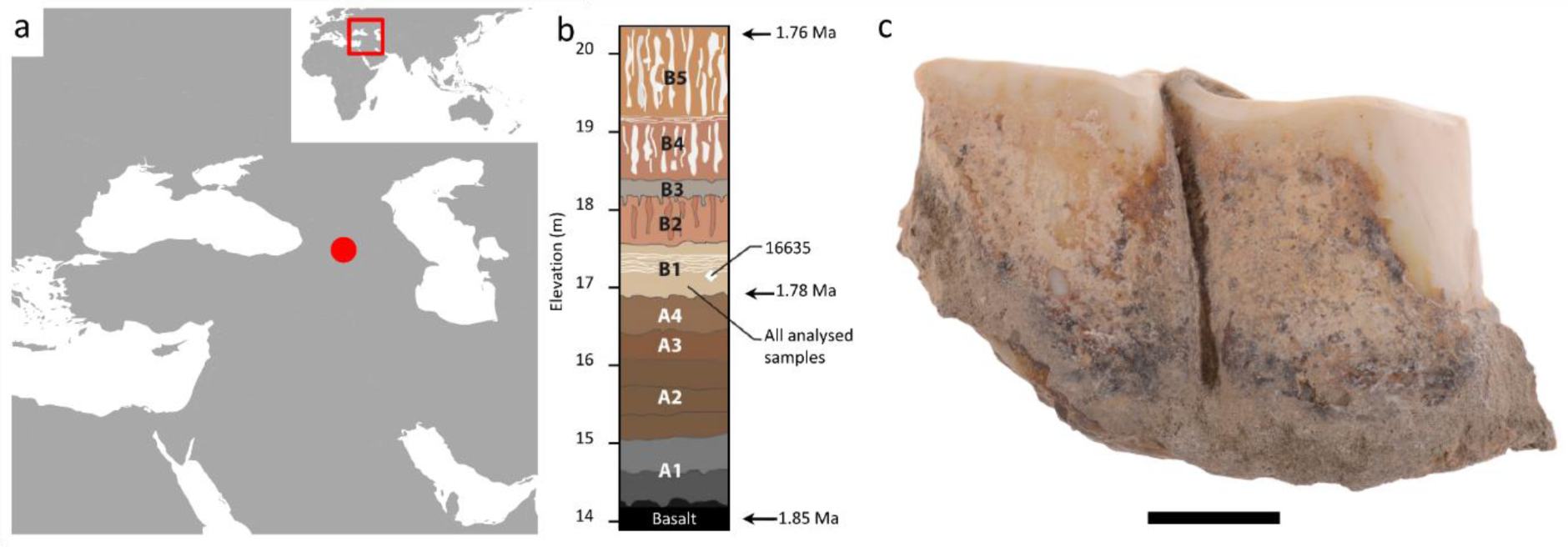
Dmanisi location, stratigraphy, and rhinoceros sample 16635. **a)** Geographic location of Dmanisi in the South Caucasus. **b)** Generalized stratigraphic profile indicating origin of the analysed specimens, recovered in layer B1 and dated to between 1.76 and 1.78 Ma. **c)** Isolated left lower molar (m1 or m2; GNM Dm.5/157-16635) of *Stephanorhinus* ex gr. *etruscus*-*hundsheimensis*, from Dmanisi (labial view). Scale bar: 1 cm.

The geology of the Dmanisi deposits provides an ideal context for the preservation of faunal materials. The primary deposits at Dmanisi are aeolian, providing for rapid, gentle burial in fine-grained, calcareous sediments. We collected 23 bone, dentine, and dental enamel specimens of large mammals (Tab. 1) from multiple excavation units within stratum B1 (Fig. 1b, Fig. 2, Tab. 1). This is an ashfall deposit that contains thousands of faunal remains, as well as all hominin fossils, in different geomorphic contexts including pipes, shallow gullies and carnivore dens. All of these are firmly dated between 1.85-1.76 Ma^18^. High-resolution tandem MS was used to confidently sequence ancient protein residues from the set of faunal remains, after digestion-free demineralisation in acid (see Methods). Ancient DNA analysis was unsuccessfully attempted on a subset of five bone and dentine specimens (see Methods).

**Table 1.** Fossil specimens selected for ancient protein and DNA extraction. For each specimen, the Centre for GeoGenetics (CGG) reference number and the Georgian National Museum (GNM) specimen field number are reported. *or the narrowest possible taxonomic identification achievable using traditional comparative anatomy methods.

| n. | CGG reference number | GNM specimen field number | Year of finding | Anatomical identification | Order | Family | Species* |
| --- | --- | --- | --- | --- | --- | --- | --- |
| 1 | CGG_1_016486 | Dm.bXI.sqA6.V._ | 1984 | P4 sin. | Canivora | Canidae | *Canis etruscus* |
| 2 | CGG_1_016626 | Dm.6/154.2/4.A4.17 | 2014 | tibia sin. | Artiodactyla | Indet. | Indet. |
| 3 | CGG_1_016628 | Dm.7/154.2.A2.27 | 2014 | mc III&IV dex. | Artiodactyla | Cervidae | Tribe Megacerini |
| 4 | CGG_1_016629 | Dm.5/154.3.A4.32 | 2014 | hemimandible sin. with dp2, dp3, dp4, m1 | Artiodactyla | Cervidae | Tribe Megacerini |
| 5 | CGG_1_016630 | Dm.6/151.4.A4.12 | 2014 | hemimandible dex. with p2-m3 | Artiodactyla | Cervidae | *Pseudodama nestii* |
| 6 | CGG_1_016631 | Dm.69/64.3.B1.53 | 2014 | maxilla sin. with P3 | Artiodactyla | Cervidae | Tribe Megacerini |
| 7 | CGG_1_016632 | Dm.5/154.2.A4.38 | 2014 | i3 dex. | Perissodactyla | Equidae | *Equus stenoris* |
| 8 | CGG_1_016633 | Dm.5/153.3.A2.33 | 2014 | mc III & mc II sin. | Perissodactyla | Equidae | *Equus stenoris* |
| 9 | CGG_1_016634 | Dm.7/151.2.B1/A4.1 | 2014 | m/1 or m/2 dex. | Perissodactyla | Equidae | *Equus stenoris* |
| 10 | CGG_1_016635 | Dm.5/157.profile cleaning | 2014 | m/1 sin. | Perissodactyla | Rhinocerotidae | *Stephanorhinus* sp. |
| 11 | CGG_1_016636 | Dm.6/153.1.A4.13 | 2014 | tibia dex. | Perissodactyla | Rhinocerotidae | Rhinocerotini indet. |
| 12 | CGG_1_016637 | Dm.7/154.2.A4.8 | 2014 | mt III&IV sin. | Artiodactyla | Bovidae | Tribe Ovibovini? Nemorhaedini? |
| 13 | CGG_1_016638 | Dm.5/154.1.B1.1 | 2014 | hemimandible dex. with p3-m3 | Artiodactyla | Bovidae | Tribe Nemorhaedini |
| 14 | CGG_1_016639 | Dm.8/154.4.A4.22 | 2014 | maxilla dex. with P2-M2 | Artiodactyla | Bovidae | Tribe Ovibovini? Nemorhaedini? |
| 15 | CGG_1_016640 | Dm.6/151.2.A4.97 | 2014 | mt III&IV sin. | Artiodactyla | Bovidae | *Bison georgicus* |
| 16 | CGG_1_016641 | Dm.8/152.3.B1.2 | 2014 | m3 dex. | Artiodactyla | Bovidae | *Bison georgicus* |
| 17 | CGG_1_016642 | Dm.8/153.4.A4.5 | 2014 | hemimandible sin. with p1-m2 | Canivora | Canidae | *Canis etruscus* |
| 18 | CGG_1_016856 | Dm.M6/7.II.296 | 2006 | m2 sin. | Artiodactyla | Cervidae | Tribe Megacerini |
| 19 | CGG_1_016857 | Dm.bXI.profile cleaning |  | long bone fragment of a herbivore | Indet. | Indet. | Indet. |
| 20 | CGG_1_016858 | Dm.bXI.North.B1a.colleciton | 2006 | metapodium fragment | Artiodactyla | Cervidae | Tribe Megacerini |
| 21 | CGG_1_016859 | D4.collection |  | fragments of pelvis and ribs of a large mammal | Indet. | Indet. | Indet. |
| 22 | CGG_1_016860 | Dm.65/62.1.A1.collection | 2011 | P4 sin. | Artiodactyla | Cervidae | Tribe Megacerini |
| 23 | CGG_1_016861 | Dm.64/63.1.B1z.collection | 2010 | fragment of an upper tooth | Perissodactyla | Equidae | *Equus stenoris* |

**Figure 2.**
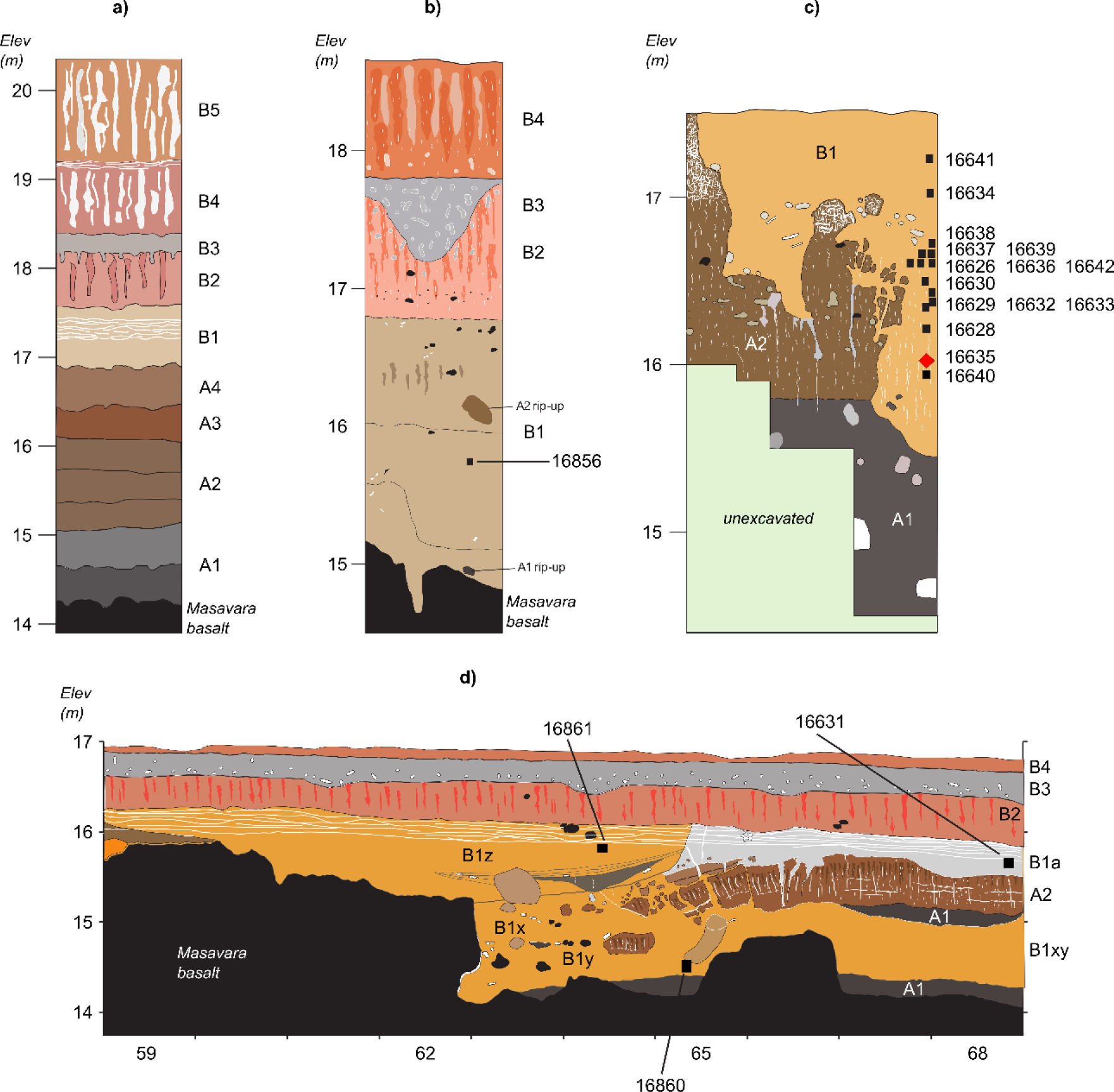
Generalized stratigraphic profiles for Dmanisi, indicating sample origins. **a)** Type section of Dmanisi in the M5 Excavation block. **b)** Stratigraphic profile of excavation area M6. M6 preserves a larger gully associated with the pipe-gully phase of stratigraphic-geomorphic development in Stratum B1. The thickness of Stratum B1 gully fill extends to the basalt surface, but includes “rip-ups” of Strata A1 and A2, showing that B1 deposits post-date Stratum A. **c)** Stratigraphic section of excavation area M17. Here, Stratum B1 was deposited after erosion of Stratum A deposits. The stratigraphic position of the *Stephanorhinus* sample 16635 is highlighted with a red diamond. The Masavara basalt is ca. 50 cm below the base of the shown profile. **d)** Northern section of Block 2. Following collapse of a pipe and erosion to the basalt, the deeper part of this area was filled with local gully fill of Stratum B1/x/y/z. Note the uniform burial of all Stratum B1 deposits by Strata B2-B4. Sampled specimens are indicated by five-digit numbers (Tab. 1). Note differences in y-axis for elevation. Five additional samples were studied from excavation area R11, stratigraphic unit B1, not shown in a stratigraphic profiles here.

While the recovery of proteins from bone and dentine specimens was sporadic and limited to collagen fragments, the analysis of dental enamel consistently returned sequences from most of its proteome, with occasional detection of multiple isoforms of the same protein^20^ (Tab. 2, Fig. 3). The small proteome^21^ of mature dental enamel consists of structural enamel proteins, i.e. amelogenin (*AMELX*), enamelin (*ENAM*), amelotin (*AMTN*), and ameloblastin (*AMBN*), and enamel-specific proteases secreted during amelogenesis, i.e. matrix metalloproteinase-20 (*MMP20*) and kallikrein 4 (*KLK4*). The presence of non-specific proteins, such as serum albumin (*ALB*), has also been previously reported in mature dental enamel^21,22^ (Tab. 2).

**Table 2.**
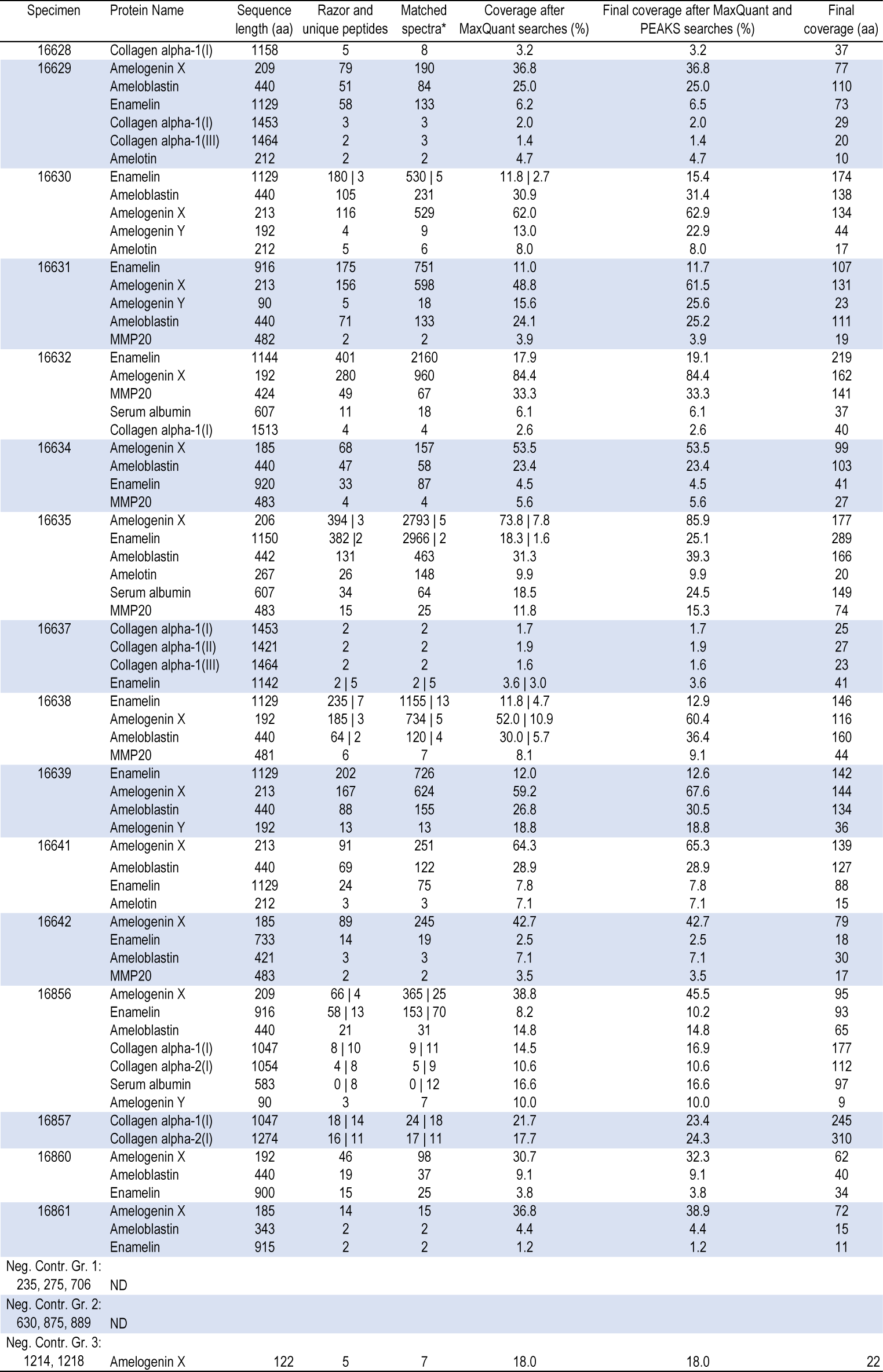
Proteome composition and coverage. In those cells reporting two values separated by the “|” symbol, the first value refers to MaxQuant (MQ) searches performed selecting unspecific digestion, while the second value refers to MQ searches performed selecting trypsin digestion. For those cells including one value only, it refers to MaxQuant (MQ) searches performed selecting unspecific digestion. Final amino acid coverage, incorporating both MQ and PEAKS searches, is reported in the last column. *supporting all peptides.

**Figure 3.**
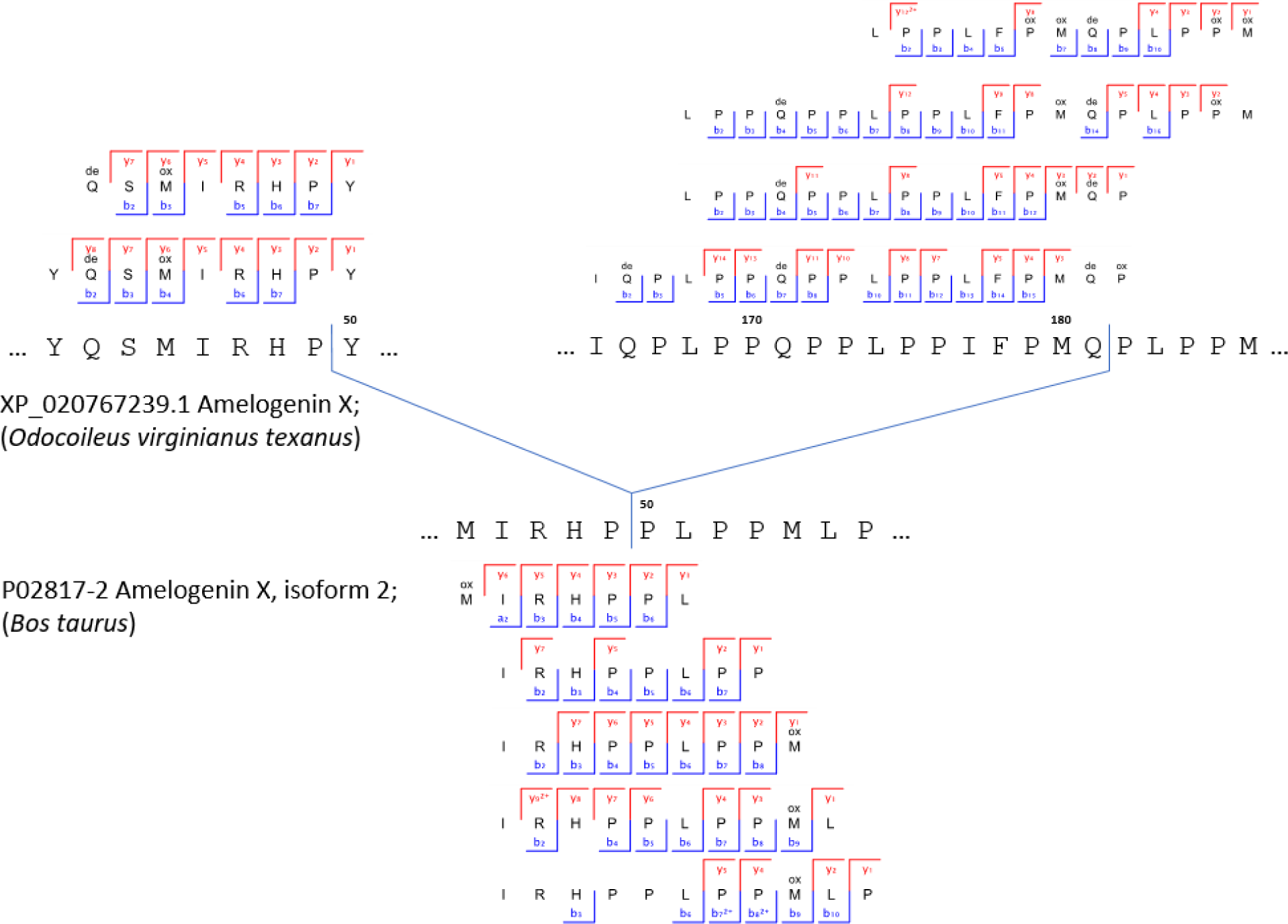
Peptide and ion fragment coverage of amelogenin X (AMELX) isoforms 1 and 2 from specimen 16856 (Cervidae). Peptides specific for amelogenin X (AMELX) isoforms 1 and 2 appear in the upper and lower parts of the figure respectively. No amelogenin X isoform 2 is currently reported in public databases for the Cervidae group. Accordingly, the amelogenin X isoform 2-specific peptides were identified by MaxQuant spectral matching against bovine (*Bos Taurus*) amelogenin X isoform 2 (UniProt accession number P02817-2). Amelogenin X isoform 2, also known as leucine-rich amelogenin peptide (LRAP), is a naturally occurring alternative Amelogenin X isoform from the translation product of an alternatively spliced transcript.

Multiple lines of evidence support the authenticity and the endogenous origin of the sequences recovered. There is full correspondence between the source material and the composition of the proteome recovered. Dental enamel proteins are extremely tissue-specific and confined to the dental enamel mineral matrix^21^. The amino acid composition of the intra-crystalline protein fraction, measured by chiral amino acid racemisation analysis, indicates that the dental enamel has behaved as a closed system, unaffected by amino acid and protein residues exchange with the burial environment (Fig. 4). The measured rate of asparagine and glutamine deamidation, a spontaneous form of hydrolytic damage consistently observed in ancient samples^23^, is particularly high, in some cases close to 100%, in full agreement with the age of the specimens investigated. (Fig. 2a). Other forms of non-enzymatic modifications are also present. Tyrosine (Y) experienced mono-and di-oxidation while tryptophan (W) was extensively converted into multiple oxidation products. (Fig. 5b). Oxidative degradation of histidine (H) and conversion of arginine (R) leading to ornithine accumulation were also observed. These modifications are absent, or much less frequent, in a medieval ovicaprine dental enamel control sample, further confirming the authenticity of the sequences reconstructed. Similarly, unlike in the control, the peptide length distribution in the Dmanisi dataset is dominated by short overlapping fragments, generated by advanced, diagenetically-induced, terminal hydrolysis (Fig. 5c and d).

**Figure 4.**
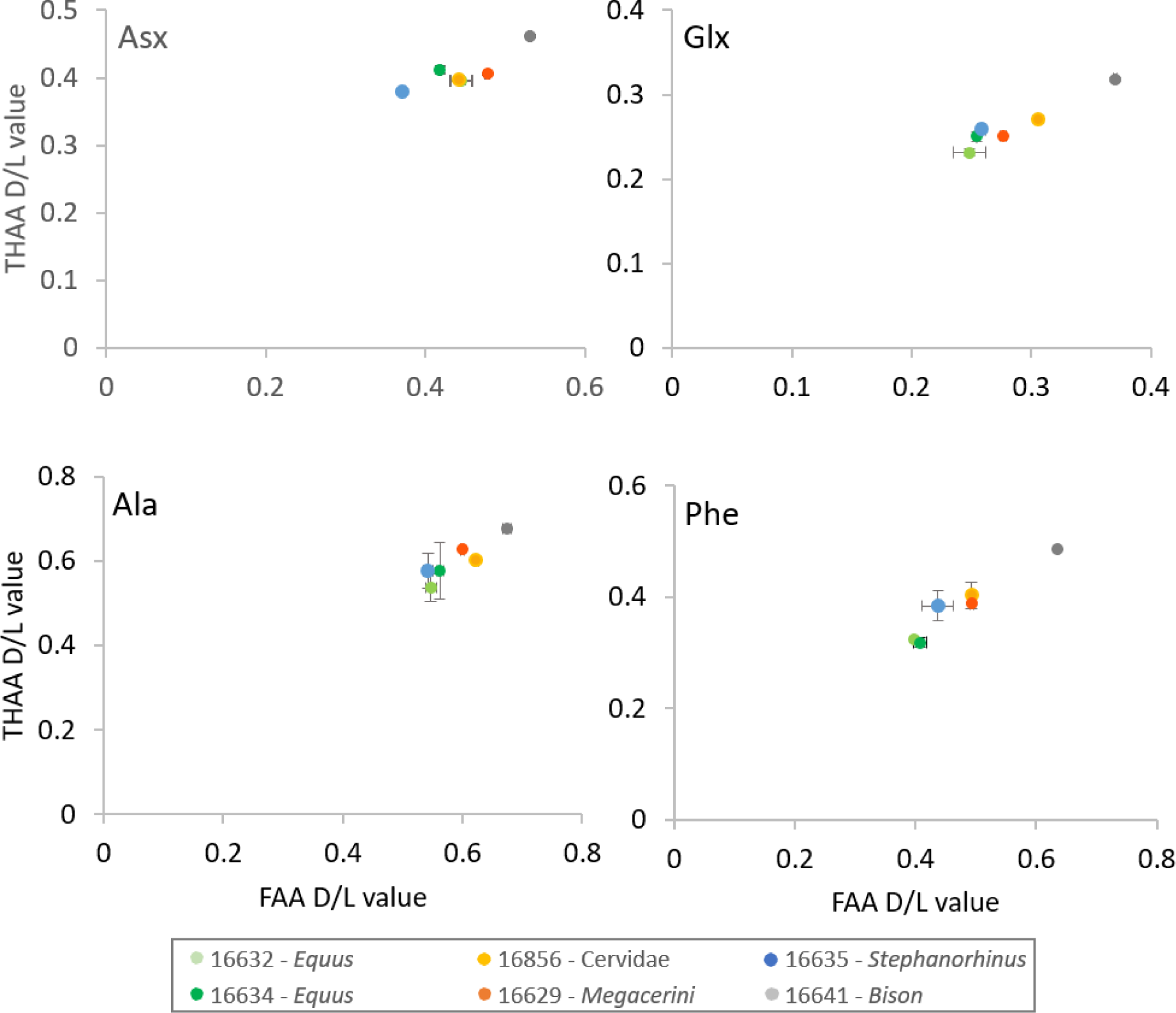
Amino Acid Racemisation. Extent of intra-crystalline racemization for four amino acids (Asx, Glx, Ala and Phe). Error bars indicate one standard deviation based on preparative replicates (n=2). “Free” amino acids (FAA) on the x-axis, “total hydrolysable” amino acids (THAA) on the y-axis. Note differences in axes for the four separate amino acids.

**Figure 5.**
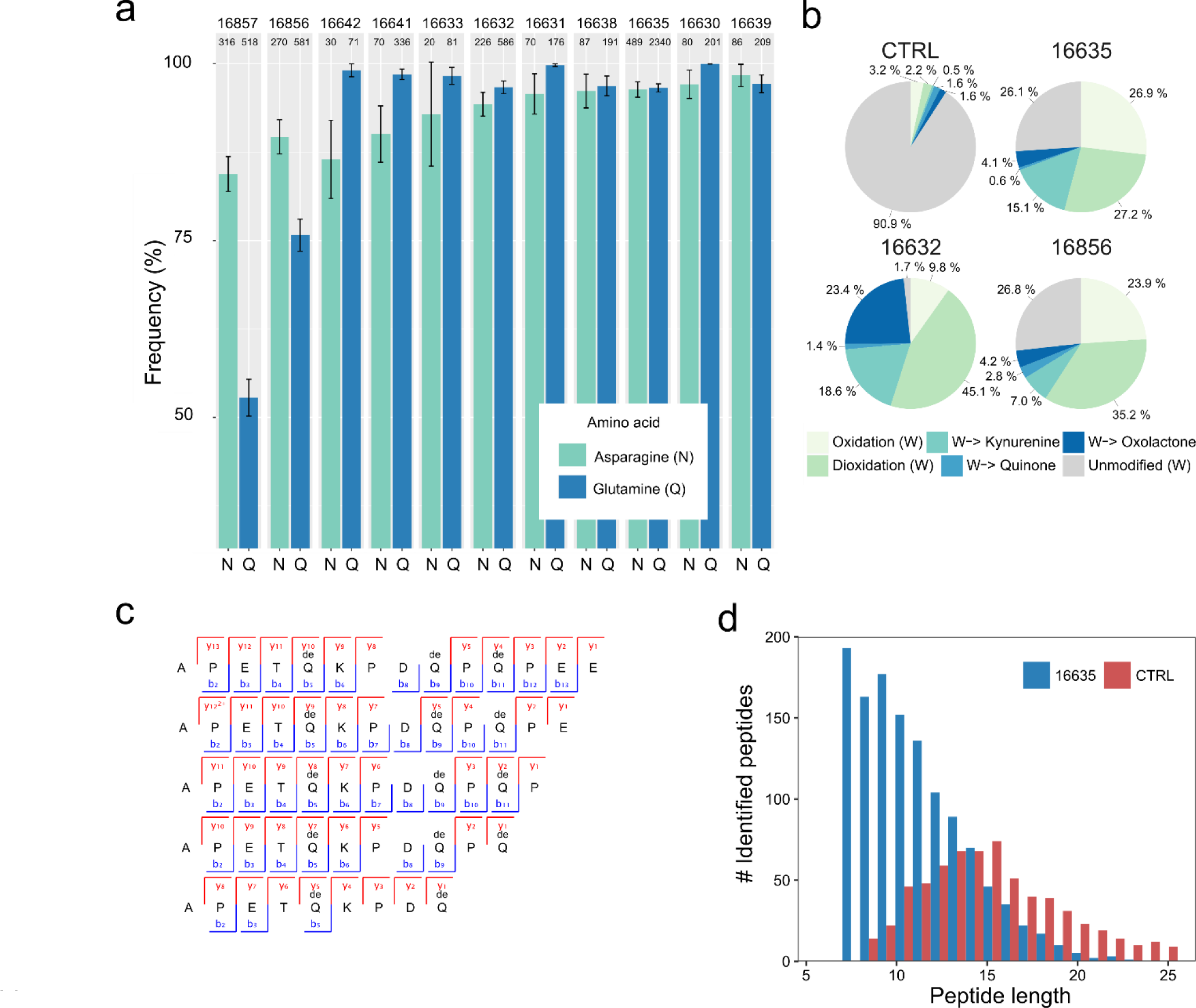
Enamel proteome degradation. **a)** Deamidation of asparagine (N) and glutamine (Q) amino acids. Error bars indicate confidence interval around 1000 bootstrap replicates. Numeric sample identifiers are shown at the very top, while the number of peptides used for the calculation are indicated for each bar. **b)** Extent of tryptophan (W) oxidation leading to several diagenetic products, measured as relative spectral counts. **c)** Peptide alignment (positions 124-137, enamelin) for acid demineralisation without enzymatic digestion. **d)** Barplot of peptide length distribution of Pleistocene *Stephanorhinus* ex gr. *etruscus*-*hundsheimensis* (16635) and Medieval (CTRL) undigested ovicaprine dental enamel proteomes, extracted and analysed in an identical manner.

Lastly, we confidently detect phosphorylation (Fig. 6 and Fig. 7), a tightly regulated physiological post-translational modification (PTM) occurring *in vivo*. Recently observed in ancient bone^24^, phosphorylation is known to be a stable PTM^25^ present in dental enamel proteins^26,27^. Altogether, these observations demonstrate, beyond reasonable doubt, that the heavily diagenetically modified dental enamel proteome retrieved from the ˜1.77 Ma old Dmanisi faunal material is endogenous and almost complete.

**Figure 6.**
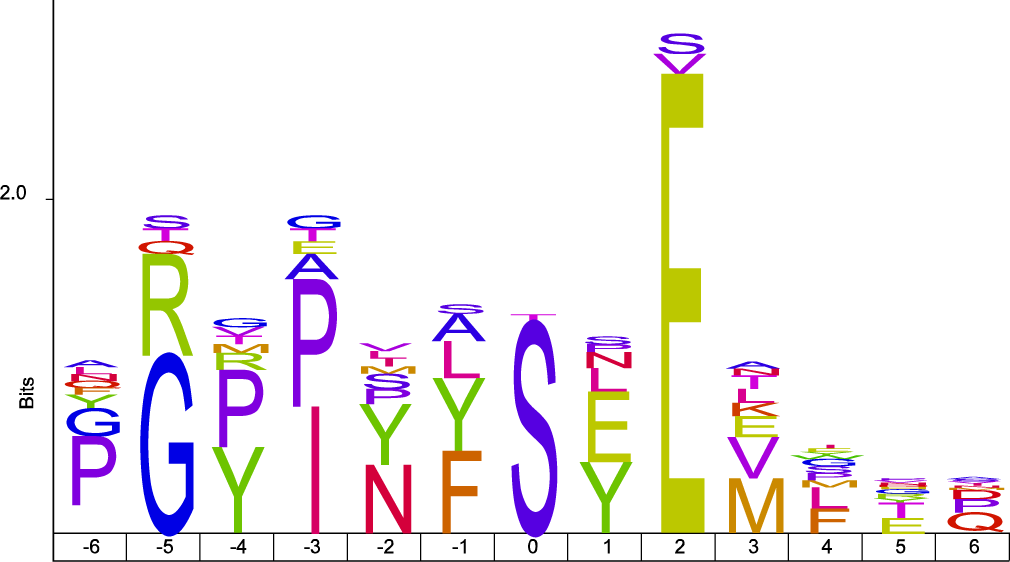
Sequence motif analysis of ancient enamel proteome phosphorylation. The identified S-x-E/phS motif is recognised by the secreted kinases of the Fam20C family, which are dedicated to the phosphorylation of extracellular proteins and involved in regulation of biomineralization^26^. See Fig. 7 for spectral examples of both S-x-E and S-x-phS phosphorylated motifs.

**Figure 7.**
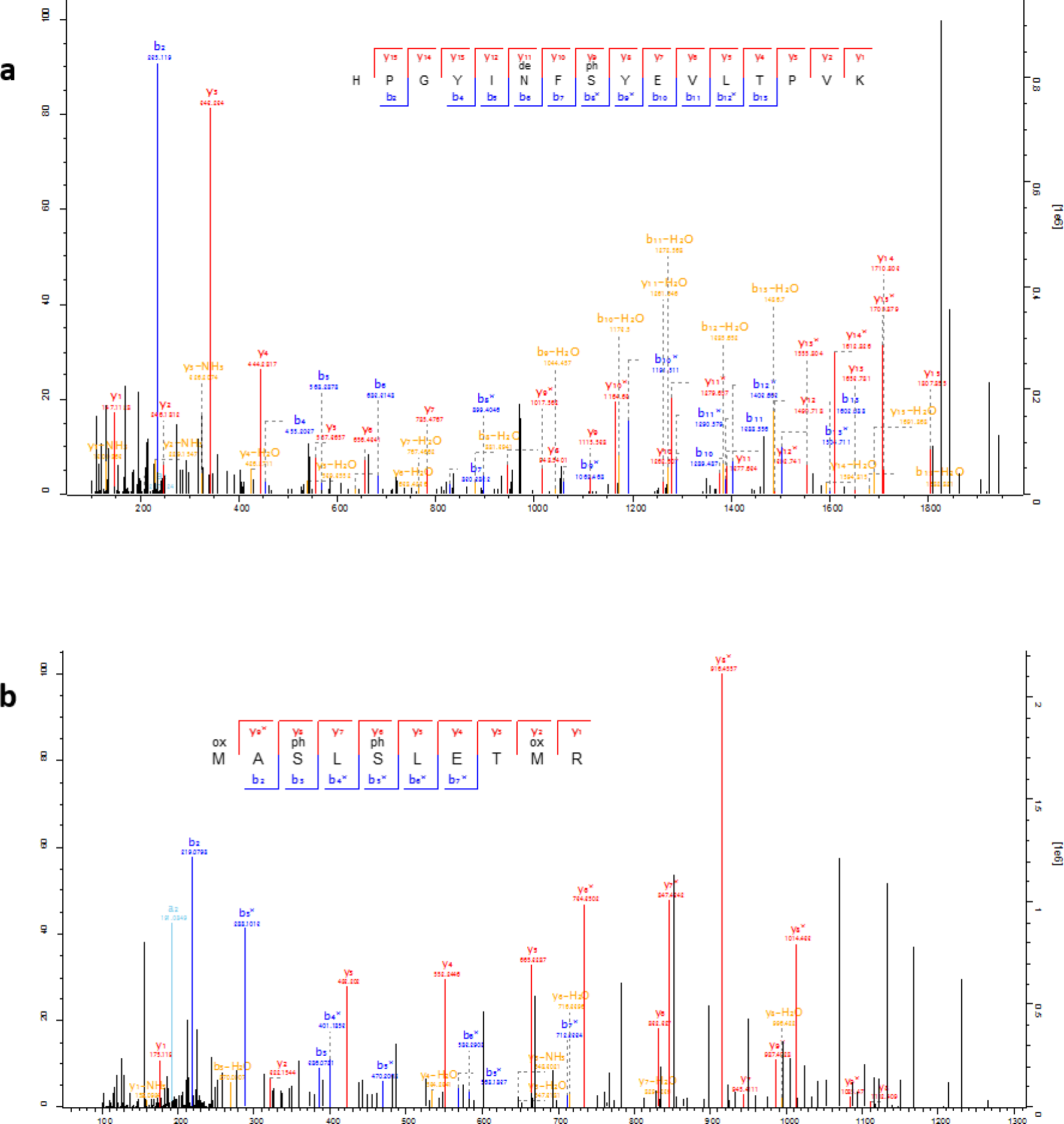
Ancient enamel proteome phosphorylation. Annotated example spectra including phosphorylated serines (phS) in the S-x-E motif **(a**; AMEL), and in the S-x-phS motif (**b;** AMBN), as well as deamidated asparagine (deN). Icelogo analysis of all phosphorylated amino acids indicates the majority derive from Fam20C kinase activity with a specificity for the phosphorylation of S-x-E or S-x-phS motifs (see Fig. 6).

Next, we used the palaeoproteomic sequence information to improve taxonomic assignment and achieve sex attribution for some of the Dmanisi faunal remains. For example, the bone specimen 16857, described morphologically as an “undetermined herbivore”, could be assigned to the Bovidae family based on COL1 sequences (Fig. 8). In addition, confident identification of peptides specific for the isoform Y of amelogenin, coded on the non-recombinant portion of the Y chromosome, indicates that four tooth specimens, namely 16630, 16631, 16639, and 16856, belonged to male individuals^22^ (Fig. 9a-d).

**Figure 8.**
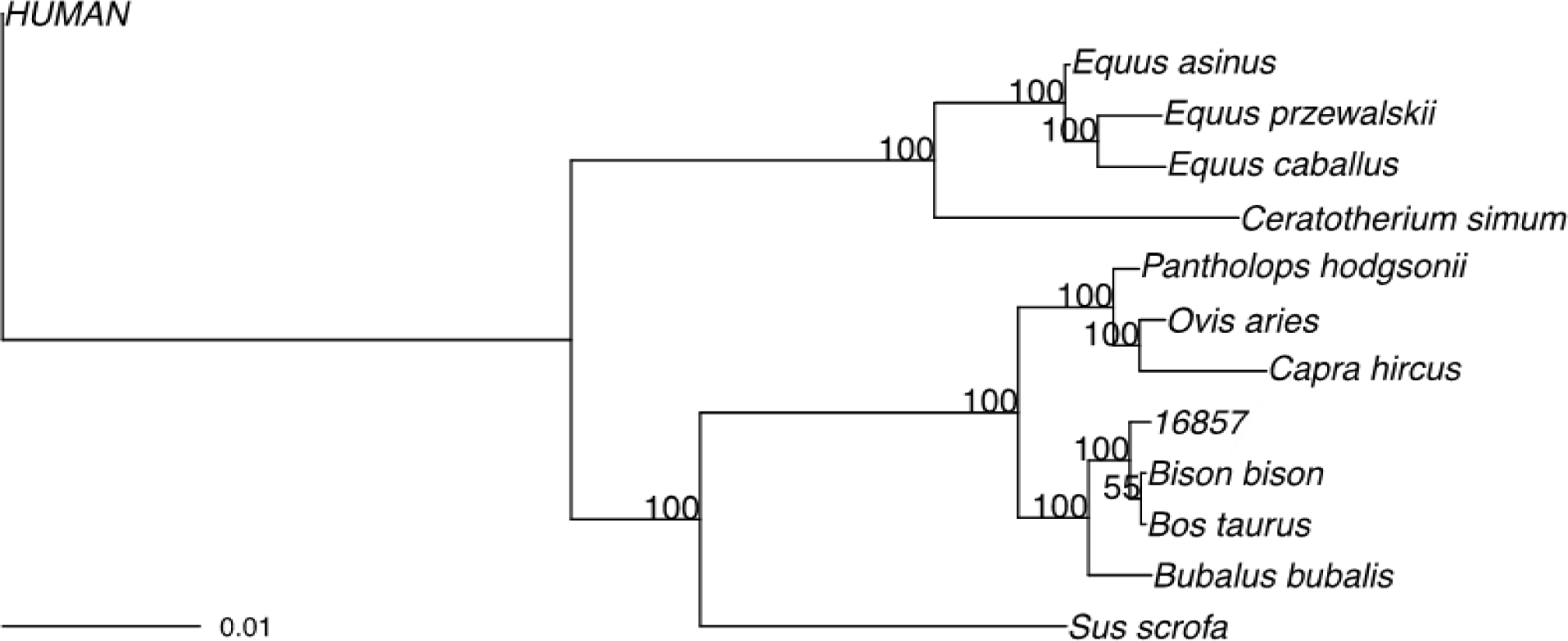
Phylogenetic relationships between the comparative reference dataset and sample 16857. Consensus tree from Bayesian inference. The posterior probability of each bipartition is shown as a percentage to the left of each node. For all panels, we show a scale for estimated branch lengths.

**Figure 9.**
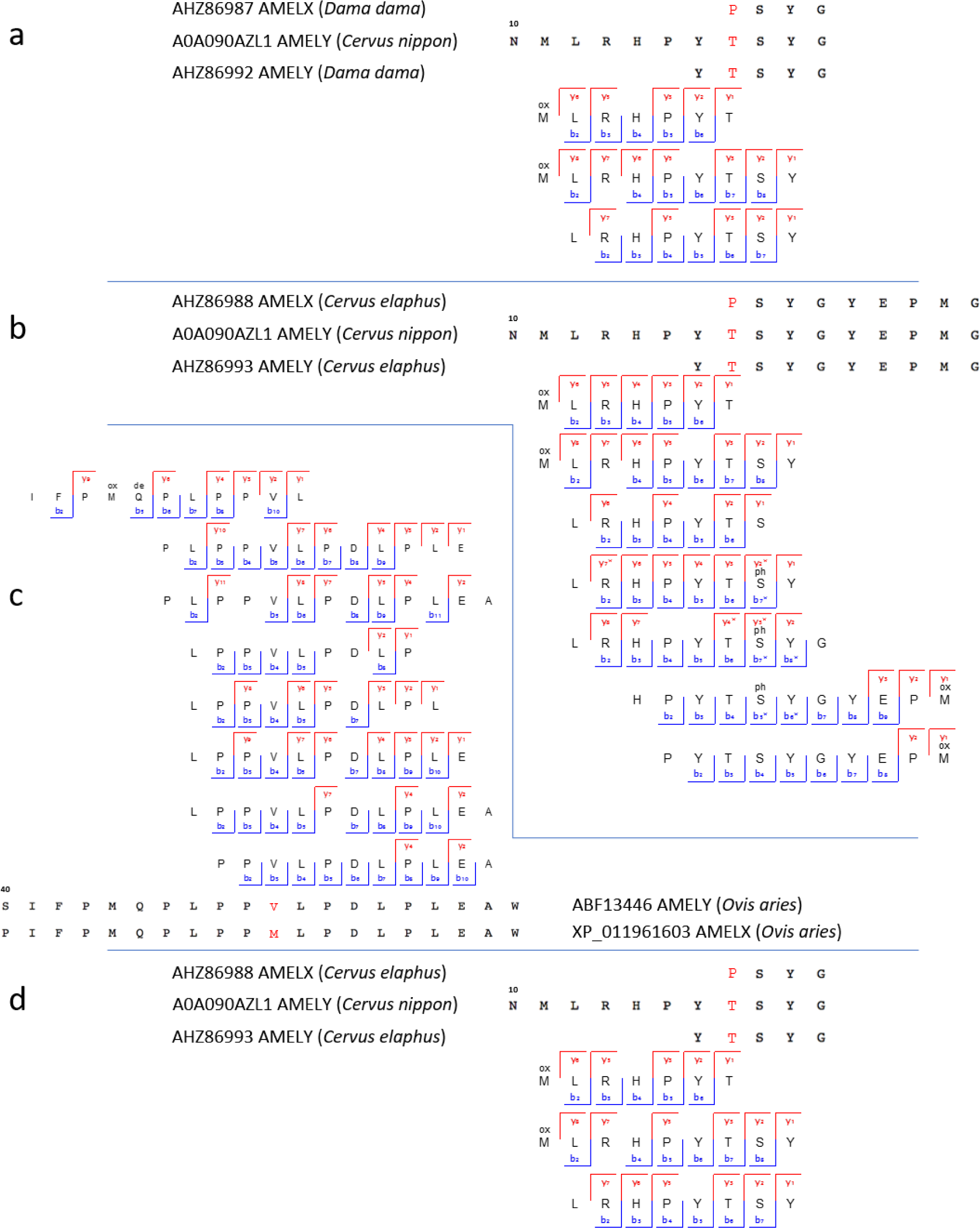
Amelogenin Y-specific matches. **a)** Sample 16630, Cervidae. **b)** Sample 16631, Cervidae. **c)** Sample 16639, Bovidae. **d)** Sample 16856, Cervidae. Note the presence of deamidated glutamines (deQ) and asparagines (deN), oxidated methionines (oxM), and phosphorylated serines (phS) in several of the indicated y- and b-ion series.

An enamel fragment, from the lower molar of a *Stephanorhinus* ex gr. *etruscus-hundsheimensis* (16635, Fig. 1c), returned the highest proteomic sequence coverage, encompassing a total of 875 amino acids, across 987 peptides (6 proteins). Following alignment of the enamel protein sequences retrieved from 16635 against their homologues from all the extant rhinoceros species, plus the extinct woolly rhinoceros (†*Coelodonta antiquitatis*) and Merck’s rhinoceros (†*Stephanorhinus kirchbergensis*), phylogenetic reconstructions place the Dmanisi specimen closer to the extinct woolly and Merck’s rhinoceroses than to the extant Sumatran rhinoceros (*Dicerorhinus sumatrensis*), as an early divergent sister lineage (Fig. 10).

**Figure 10.**
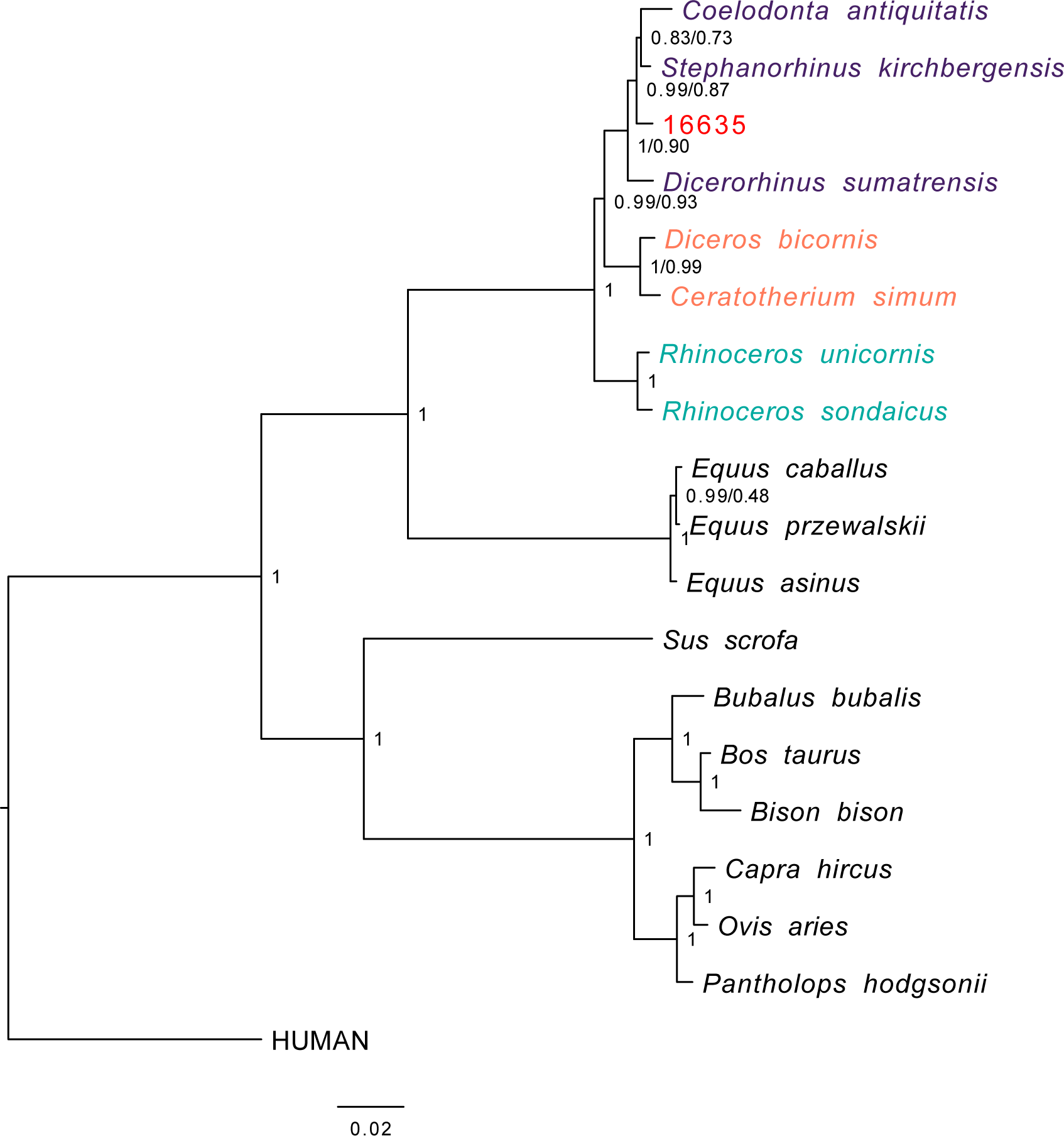
Phylogenetic relationships between the comparative enamel proteome dataset and specimen 16635 (*Stephanorhinus* ex gr. *etruscus-hundsheimensis*). Consensus tree from Bayesian inference on the concatenated alignment of six enamel proteins and using *Homo sapiens* as an outgroup. For each bipartition, we show the posterior probability obtained from the Bayesian inference. Additionally, for bipartitions where the Bayesian and the Maximum-likelihood inference support are different, we show (right) the support obtained in the latter. Scale indicates estimated branch lengths. Colours indicate the three main rhinoceros clades: Sumatran-extinct (purple), African (orange) and Indian-Javan (green), as well as the specimen 16635 (red).

**Figure 11.**
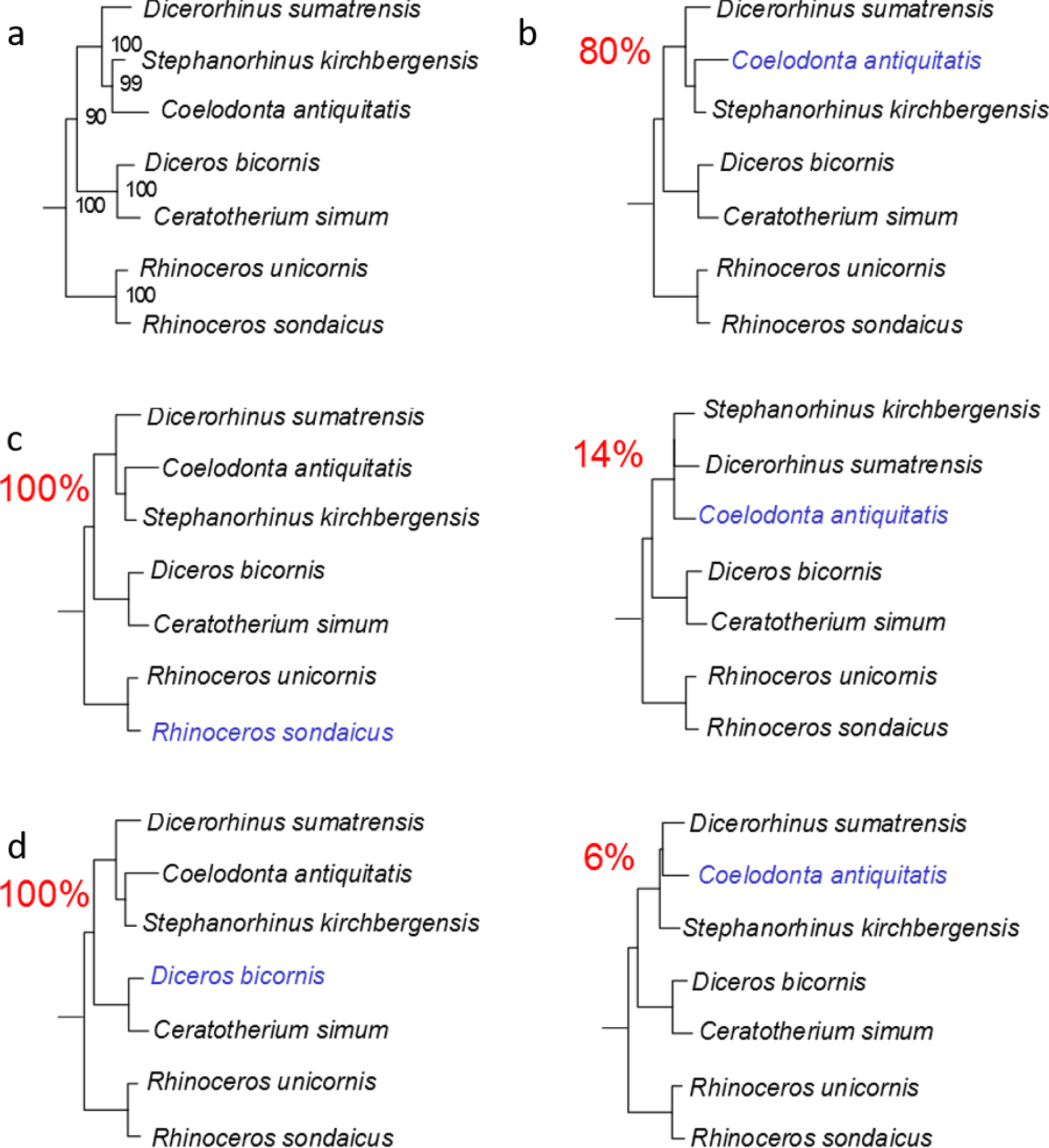
Effect of the missingness in the tree topology. **a)** Maximum-likelihood phylogeny obtained using PhyML and the protein alignment excluding the ancient Dmanisi rhinoceros. **b)** Topologies obtained from 100 random replicates of the Woolly rhinoceros (*Coelodonta antiquitatis*). Each replicate was added a similar amount of missing sites as in the Dmanisi sample (72.4% missingness). The percentage shown for each topology indicates the number of replicates in which that particular topology was recovered. **c)** Similar to **b**, but for the Javan rhinoceros (*Rhinoceros sondaicus*). **d)** Similar to **b**, but for the black rhinoceros (*Diceros bicornis*).

Our phylogenetic reconstruction confidently recovers the expected differentiation of the *Rhinoceros* genus from other genera considered, in agreement with previous cladistic^28^ and genetic analyses^29^. This topology defines two-horned rhinoceroses as monophyletic and the one-horned condition as plesiomorphic, as previously proposed^30^. We caution, however, that the higher-level relationships we observe between the rhinoceros monophyletic clades might be affected by demographic events, such as incomplete lineage sorting^31^ and/or gene flow between groups^32^, due to the limited number of markers considered. A previous phylogenetic reconstruction, based on two collagen (*COL1α1* and *COL1α2*) partial amino acid sequences, supported a different topology, with the African clade representing an outgroup to Asian rhinoceros species^6^. Most probably, a confident and stable reconstruction of the structure of the Rhinocerotidae family needs the strong support only high-resolution whole-genome sequencing can provide. Regardless, the highly supported placement of the Dmanisi rhinoceros in the (*Stephanorhinus*, Woolly, Sumatran) clade will likely remain unaffected, should deeper phylogenetic relationships between the *Rhinoceros* genus and other family members be revised.

The phylogenetic relationships of the genus *Stephanorhinus* within the family Rhinocerotidae, as well as those of the several species recognized within this genus, are contentious. *Stephanorhinus* was initially included in the extant South-East Asian genus *Dicerorhinus* represented by the Sumatran rhinoceros species (*D. sumatrensis*)^33^. This hypothesis has been rejected and, based on morphological data, *Stephanorhinus* has been identified as a sister taxon of the woolly rhinoceros^34^. Furthermore, ancient DNA analysis supports a sister relationship between the woolly rhinoceros and *D. sumatrensis* ^5,35,36^.

Recently, MS-based sequencing of collagen type I from a Middle Pleistocene European *Stephanorhinus* sp. specimen, ˜320 ka (thousand years) old, was not able to resolve the relationships between *Stephanorhinus, Coelodonta* and *Dicerorhinus*^6^. Instead, the complete mitochondrial sequence of a terminal, 45-70 ka old, Siberian *S. kirchbergensis* specimen placed this species closer to *Coelodonta*, with *D. sumatrensis* as a sister branch^7^. Our results confirm the latter reconstruction. As the *Stephanorhinus* ex gr. *etruscus-hundsheimensis* sequences from Dmanisi branch off basal to the common ancestor of the woolly and Merck’s rhinoceroses, these two species most likely derived from an early *Stephanorhinus* lineage expanding eastward from western Eurasia. Throughout the Plio-Pleistocene, *Coelodonta* adapted to continental and later cold-climate habitats in central Asia. Its earliest representative, *C. thibetana,* displayed some clear *Stephanorhinus*-like anatomical features^34^. The presence in eastern Europe and Anatolia of the genus *Stephanorhinus*^35^ is documented at least since the late Miocene, and the Dmanisi specimen most likely represents an Early Pleistocene descendent of the Western-Eurasian branch of this genus.

Ultimately, our phylogenetic reconstructions show that, as currently defined, the genus *Stephanorhinus* is paraphyletic, in line with previous conclusions^37^ based on morphological characters and the palaeobiogeographic fossil distribution. Accordingly, a systematic revision of the genera *Stephanorhinus* and *Coelodonta*, as well as their closest relatives, is needed.

In this study, we show that enamel proteome sequencing can overcome the time limits of ancient DNA preservation and the reduced phylogenetic content of COL1 sequences. Dental enamel proteomic sequences can be used to study evolutionary process that occurred in the Early Pleistocene. This posits dental enamel as the material of choice for deep-time palaeoproteomic analysis. Given the abundance of teeth in the palaeontological record, the approach presented here holds the potential to address a wide range of questions pertaining to the Early and Middle Pleistocene evolutionary history of a large number of mammals, including hominins, at least in temperate climates.

## METHODS

### Dmanisi & sample selection

Dmanisi is located about 65 km southwest of the capital city of Tbilisi in the Kvemo Kartli region of Georgia, at an elevation of 910 m MSL (Lat: 41° 20’ N, Lon: 44° 20’ E)^8,19^. The 23 fossil specimens we analysed were retrieved from stratum B1, in excavation blocks M17, M6, block 2, and area R11 (Tab. 1 and Fig. 2). Stratum B deposits date between 1.78 Ma and 1.76 Ma^18^. All the analysed specimens were collected between 1984 and 2014 and their taxonomic identification was based on traditional comparative anatomy.

After the sample preparation and data acquisition for all the Dmanisi specimens was concluded, we applied the whole experimental procedure to a medieval ovicaprine (sheep/goat) dental enamel specimen that was used as control. For this sample, we used extraction protocol “C”, and generated tandem MS data using a Q Exactive HF mass spectrometer (Thermo Fisher Scientific). The data were searched against the goat proteome, downloaded from the NCBI Reference Sequence Database (RefSeq) archive^38^ on 31^st^ May 2017. The ovicaprine specimen was found at the “Hotel Skandinavia” site in the city of Århus, Denmark and was stored at the Natural History Museum of Denmark.

### Biomolecular preservation

We assessed the potential of ancient protein preservation prior to proteomic analysis by measuring the extent of amino acid racemisation in a subset of samples (6/23)^39^. Enamel chips were powdered, and two subsamples per specimen were subject to analysis of their free (FAA) and total hydrolysable (THAA) amino acid fractions. Samples were analysed in duplicate by RP-HPLC, with standards and blanks run alongside each one of them. The D/L values of aspartic acid/asparagine, glutamic acid/glutamine, phenylalanine and alanine (D/L Asx, Glx, Phe, Ala) were assessed (Fig. 4) to provide an overall estimate of intra-crystalline protein decomposition (IcPD).

### PROTEOMICS

All the sample preparation procedures for palaeoproteomic analysis were conducted in laboratories dedicated to the analysis of ancient DNA and ancient proteins in clean rooms fitted with filtered ventilation and positive pressure, in line with recent recommendations for ancient protein analysis^40^. A mock “extraction blank”, containing no starting material, was prepared, processed and analysed together with each batch of ancient samples.

### Sample preparation

The external surface of bone and dentine samples was gently removed, and the remaining material was subsequently powdered. Enamel fragments, occasionally mixed with small amounts of dentine, were removed from teeth with a cutting disc and subsequently crushed into a rough powder. Ancient protein residues were extracted from approximately 180-220 mg of mineralised material, unless otherwise specified, using three different extraction protocols, hereafter referred to as “**A**”, “**B**” and “**C**”:

#### EXTRACTION PROTOCOL A - FASP

Tryptic peptides were generated using a filter-aided sample preparation (FASP) approach^41^, as previously performed on ancient samples^42^.

#### EXTRACTION PROTOCOL B - GuHCl solution and digestion

Bone or dentine powder was demineralised in 1 mL 0.5 M EDTA pH 8.0. After removal of the supernatant, all demineralised pellets were re-suspended in a 300 µL solution containing 2 M guanidine hydrochloride (GuHCl, Thermo Scientific), 100 mM Tris pH 8.0, 20 mM 2-Chloroacetamide (CAA), 10 mM Tris (2-carboxyethyl)phosphine (TCEP) in ultrapure H_2_O^43,44^. A total of 0.2 µg of mass spectrometry-grade rLysC (Promega P/N V1671) enzyme was added before the samples were incubated for 3-4 hours at 37°C with agitation. Samples and negative controls were subsequently diluted to 0.6 M GuHCl, and 0.8 µg of mass spectrometry-grade Trypsin (Promega P/N V5111) was added. The entire amount of extracted proteins was digested. Next, samples and negative controls were incubated overnight under mechanical agitation at 37°C. On the following day, samples were acidified, and the tryptic peptides were immobilised on Stage-Tips, as previously described^45^.

#### EXTRACTION PROTOCOL C - digestion-free acid demineralisation

Dental enamel powder was demineralised in 1.2 M HCl at room temperature, after which the solubilised protein residues were directly cleaned and concentrated on Stage-Tips, as described above. The sample prepared on Stage-Tip “#1217” was processed with 10% TFA instead of 1.2 M HCl. All the other parameters and procedures were identical to those used for all the other samples extracted with protocol “**C**”.

### Tandem mass spectrometry

Different sets of samples were analysed by nanoflow liquid chromatography coupled to tandem mass spectrometry (nanoLC-MS/MS) on an EASY-nLC™ 1000 or 1200 system connected to a Q-Exactive, a Q-Exactive Plus, or to a Q-Exactive HF (Thermo Scientific, Bremen, Germany) mass spectrometer. Before and after each MS/MS run measuring ancient or extraction blank samples, two successive MS/MS run were included in the sample queue in order to prevent carryover contamination between the samples. These consisted, first, of a MS/MS run (“MS/MS blank” run) with an injection exclusively of the buffer used to re-suspend the samples (0.1% TFA, 5% ACN), followed by a second MS/MS run (“MS/MS wash” run) with no injection.

### Data analysis

Raw data files generated during MS/MS spectral acquisition were searched using MaxQuant^46^, version 1.5.3.30, and PEAKS^47^, version 7.5. A two-stage peptide-spectrum matching approach was adopted. Raw files were initially searched against a target/reverse database of collagen and enamel proteins retrieved from the UniProt and NCBI Reference Sequence Database (RefSeq) archives^38,48^, taxonomically restricted to mammalian species. A database of partial “COL1A1” and “COL1A2” sequences from cervid species^13^ was also included. The results from the preliminary analysis were used for a first, provisional reconstruction of protein sequences.

For specimens whose dataset resulted in a narrower, though not fully resolved, initial taxonomic placement, a second MaxQuant search (MQ2) was performed using a new protein database taxonomically restricted to the “order” taxonomic rank as determined after MQ1. For the MQ2 matching of the MS/MS spectra from specimen 16635, partial sequences of serum albumin and enamel proteins from Sumatran (*Dicerorhinus sumatrensis*), Javan (*Rhinoceros sondaicus*), Indian (*Rhinoceros unicornis*), woolly (*Coelodonta antiquitatis*), Mercks (*Stephanorhinus kirchbergensis*), and Black rhinoceros (*Diceros bicornis*), were also added to the protein database. All the protein sequences from these species were reconstructed from draft genomes for each species (Dalen and Gilbert, unpublished data).

For each MaxQuant and PEAKS search, enzymatic digestion was set to “unspecific” and the following variable modifications were included: oxidation (M), deamidation (NQ), N-term Pyro-Glu (Q), N-term Pyro-Glu (E), hydroxylation (P), phosphorylation (S). The error tolerance was set to 5 ppm for the precursor and to 20 ppm, or 0.05 Da, for the fragment ions in MaxQuant and PEAKS respectively. For searches of data generated from sample fractions partially or exclusively digested with trypsin, another MaxQuant and PEAKS search was conducted using the “enzyme” parameter set to “Trypsin/P”. Carbamidomethylation (C) was set: (i) as a fixed modification, for searches of data generated from sets of sample fractions exclusively digested with trypsin, or (ii) as a variable modification, for searches of data generated from sets of sample fractions partially digested with trypsin. For searches of data generated exclusively from undigested sample fractions, carbamidomethylation (C) was not included as a modification, neither fixed nor variable.

The datasets re-analysed with MQ2 search, were also processed with the PEAKS software using the entire workflow (PEAKS *de novo* to PEAKS SPIDER) in order to detect hitherto unreported single amino acid polymorphisms (SAPs). Any amino acid substitution detected by the “SPIDER” homology search algorithm was validated by repeating the MaxQuant search (MQ3). In MQ3, the protein database used for MQ2 was modified to include the amino acid substitutions detected by the “SPIDER” algorithm.

### Ancient protein sequence reconstruction

The peptide sequences confidently identified by the MQ1, MQ2, MQ3 were aligned using the software Geneious^49^ (v. 5.4.4, substitution matrix BLOSUM62, gap open penalty 12 and gap extension penalty). The peptide sequences confidently identified by the PEAKS searches were aligned using an in-house R-script. A consensus sequence for each protein from each specimen was generated in FASTA format, without filtering on depth of coverage. Amino acid positions that were not confidently reconstructed were replaced by an “X”. We took into account variable leucine/isoleucine, glutamine/glutamic acid, and asparagine/aspartic acid positions through manual interpretation of possible conflicting positions (leucine/isoleucine) and replacement of possibly deamidated positions into “X” for phylogenetically informative sites. The output of the MQ2 and 3 peptide-spectrum matching was used to extend the coverage of the ancient protein sequences initially identified in the MQ1 iteration.

### Post translational modifications

#### Deamidation

After removal of likely contaminants, the extent of glutamine and asparagine deamidation was estimated for individual specimens, by using the MaxQuant output files as previously published^44^.

#### Other spontaneous chemical modifications

Spontaneous post-translational modifications (PTMs) associated with chemical protein damage were searched using the PEAKS PTM tool and the dependent peptides search mode^50^ in MaxQuant. In the PEAKS PTM search, all modifications in the Unimod database were considered. The mass error was set to 5.0 ppm and 0.5 Da for precursor and fragment, respectively. For PEAKS, the *de novo* ALC score was set to a threshold of 15 % and the peptide hit threshold to 30. The results were filtered by an FDR of 5 %, *de novo* ALC score of 50 %, and a protein hit threshold of ≥ 20. The MaxQuant dependent peptides search was carried out with the same search settings as described above and with a dependent peptide FDR of 1 % and a mass bin size of 0.0065 Da. For validation purposes, up to 10 discovered modifications were specified as variable modifications and re-searched with MaxQuant. The peptide FDR was manually adjusted to 5 % on PSM level and the PTMs were semi-quantified by relative spectral counting.

#### Phosphorylation

Class I phosphorylation sites were selected with localisation probabilities of ≥0.98 in the Phosph(ST)Sites MaxQuant output file. Sequence windows of ±6 aa from all identified sites were compared against a background file containing all non-phosphorylated peptides using a linear kinase sequence motif enrichment analysis in IceLogo^51^.

## PHYLOGENETIC ANALYSIS

### Reference datasets

We assembled a reference dataset consisting of publicly available protein sequences from representative ungulate species belonging to the following families: Equidae, Rhinocerotidae, Suidae and Bovidae. We extended this dataset with the protein sequences from extinct and extant rhinoceros species including: the woolly rhinoceros (†*Coelodonta antiquitatis*), the Merck’s rhinoceros (†*Stephanorhinus kirchbergensis*), the Sumatran rhinoceros (*Dicerorhinus sumatrensis*), the Javan rhinoceros (*Rhinoceros sondaicus*), the Indian rhinoceros (*Rhinoceros unicornis*), and the Black rhinoceros (*Diceros bicornis*). Their corresponding protein sequences were obtained following translation of high-throughput DNA sequencing data, after filtering reads with mapping quality lower than 30 and nucleotides with base quality lower than 20, and calling the majority rule consensus sequence using ANGSD^52^ For the woolly and Merck’s rhinoceroses we excluded the first and last five nucleotides of each DNA fragment in order to minimize the effect of postmortem ancient DNA damage^53^. Each consensus sequence was formatted as a separate blast nucleotide database. We then performed a tblastn^54^ alignment using the corresponding white rhinoceros sequence as a query, favouring ungapped alignments in order to recover translated and spliced protein sequences. Resulting alignments were processed using ProSplign algorithm from the NCBI Eukaryotic Genome Annotation Pipeline^55^ to recover the spliced alignments and translated protein sequences.

### Construction of phylogenetic trees

For each specimen, multiple sequence alignments for each protein were built using mafft^56^ and concatenated onto a single alignment per specimen. These were inspected visually to correct obvious alignment mistakes, and all the isoleucine residues were substituted with leucine ones to account for indistinguishable isobaric amino acids at the positions where the ancient protein carried one of such amino acids. Based on these alignments, we inferred the phylogenetic relationship between the ancient samples and the species included in the reference dataset by using three approaches: distance-based neighbour-joining, maximum likelihood and Bayesian phylogenetic inference.

Neighbour-joining trees were built using the phangorn^57^ R package, restricting to sites covered in the ancient samples. Genetic distances were estimated using the JTT model, considering pairwise deletions. We estimated bipartition support through a non-parametric bootstrap procedure using 500 pseudoreplicates. We used PHyML 3.1^58^ for maximum likelihood inference based on the whole concatenated alignment. For likelihood computation, we used the JTT substitution model with two additional parameters for modelling rate heterogeneity and the proportion of invariant sites. Bipartition support was estimated using a non-parametric bootstrap procedure with 500 replicates. Bayesian phylogenetic inference was carried out using MrBayes 3.2.6^59^ on each concatenated alignment, partitioned per gene. While we chose the JTT substitution model in the two approaches above, we allowed the Markov chain to sample parameters for the substitution rates from a set of predetermined matrices, as well as the shape parameter of a gamma distribution for modelling across-site rate variation and the proportion of invariable sites. The MCMC algorithm was run with 4 chains for 5,000,000 cycles. Sampling was conducted every 500 cycles and the first 25% were discarded as burn-in. Convergence was assessed using Tracer v. 1.6.0, which estimated an ESS greater than 5,500 for each individual, indicating reasonable convergence for all runs.

## ANCIENT DNA ANALYSIS

The samples were processed using strict aDNA guidelines in a clean lab facility at the Centre for GeoGenetics, Natural History Museum of Denmark, University of Copenhagen. DNA extraction was attempted on five of the ancient animal samples. Powdered samples (120-140 mg) were extracted using a silica-in-solution method^10,60^. To prepare the samples for NGS sequencing, 20 μL of DNA extract was built into a blunt-end library using the NEBNext DNA Sample Prep Master Mix Set 2 (E6070) with Illumina-specific adapters. The libraries were PCR-amplified with inPE1.0 forward primers and custom-designed reverse primers with a 6-nucleotide index^61^. Two extracts (MA399 and MA2481, from specimens 16859 and 16635 respectively) yielded detectable DNA concentrations. These extracts were used to construct three individual index-barcoded libraries (MA399_L1, MA399_L2, MA2481_L1) whose amplification required a total of 30 PCR cycles in a 2-round setup (12 cycles with total library + 18 cycles with a 5 μL library aliquot from the first amplification). The libraries generated from specimen 16859 and 16635 were processed on different flow cells. They were pooled with others for sequencing on an Illumina 2000 platform (MA399_L1, MA399_L2) using 100bp single read chemistry and on an Illumina 2500 platform (MA2481_L1) using 81bp single read chemistry.

The data were base-called using the Illumina software CASAVA 1.8.2 and sequences were demultiplexed with a requirement of a full match of the six nucleotide indexes that were used. Raw reads were processed using the PALEOMIX pipeline following published guidelines^62^, mapping against the cow nuclear genome (*Bos taurus* 4.6.1, accession GCA_000003205.4), the cow mitochondrial genome (*Bos taurus*), the red deer mitochondrial genome (*Cervus elaphus*, accession AB245427.2), and the human nuclear genome (GRCh37/hg19), using BWA backtrack^63^ v0.5.10 with the seed disabled. All other parameters were set as default. PCR duplicates from mapped reads were removed using the picard tool *MarkDuplicate* [http://picard.sourceforge.net/].

## SAMPLE 16635 MORPHOLOGICAL MEASUREMENTS

We followed the methodology introduced by Guérin^33^. The maximal length of the tooth is measured with a digital calliper at the lingual side of the tooth and parallel to the occlusal surface. All measurements are given in mm.

## DATA DEPOSITION

All the mass spectrometry proteomics data have been deposited in the ProteomeXchange Consortium (http://proteomecentral.proteomexchange.org) via the PRIDE partner repository with the data set identifier PXD011008.

## ACKNOWLEDGEMENTS

We would like to thank, Kristian Murphy Gregersen, for providing the medieval control specimen, Marcus Anders Krag for the photographs used in Fig. 1c, Fedor Shidlovskiy for providing access to the Merck’s rhino sample, Beatrice Triozzi for technical help, Ashot Margaryan and Shyam Gopalakrishnan for their precious comments during data interpretation. EC and FW are supported by VILLUM Fonden (grant number 17649). EC, CK, JVO, PR and DS are supported by the Marie Skłodowska Curie European Training Network “TEMPERA” (grant number 722606). MM is supported by the University of Copenhagen KU2016 (UCPH Excellence Programme) grant and by the Danish National Research Foundation award PROTEIOS (DNRF128). Work at the Novo Nordisk Foundation Center for Protein Research is funded in part by a generous donation from the Novo Nordisk Foundation (Grant number NNF14CC0001). MTPG is supported by ERC Consolidator Grant “EXTINCTION GENOMICS” (grant number 681396). LP was supported by the EU-SYNTHESYS project (AT-TAF-2550, DE-TAF-3049, GB-TAF-2825, HU-TAF-3593, ES-TAF-2997) funded by the European Commission. LO is supported by the ERC Consolidator Grant “PEGASUS” (grant agreement No 681605). BM-N is supported by the Spanish Ministry of Sciences (grant number CGL2016-80975-P). BS, JK and PH are supported by the Gordon and Betty Moore foundation. The aDNA analysis was carried out using the HPC facilities of the University of Luxembourg.

## AUTHOR CONTRIBUTIONS

E.C., D.Lo., and E.W. designed the study. A.K.F., M.M., R.R.J.-C., M.E.A., M.D., K.P., and E.C. performed laboratory experiments. M.Bu., M.T., R.F., E.P., T.S., Y.L.C., A.Gö., S.N., P.H., J.K., I.K., Y.M., J.A., R.-D.K., G.K., B.M.-N., M.-H.S.S., S.L., M.S.V., B.S., L.D., M.T.P.G., and D.Lo., provided ancient samples or modern reference material. E.C., F.W., L.P., J.R.M., D.Ly, V.J.M.M., A.K., D.S., C.K., A.Gi., L.O., L.R., J.V.O., P.R., M.D., and K.P. performed analyses and data interpretation. E.C., F.W., J.R.M., L.P. and E.W. wrote the manuscript with contributions of all authors.

